# Immune response to spiral ganglion neuron death in rats during development and after kanamycin-induced deafening

**DOI:** 10.64898/2026.03.10.710901

**Authors:** Adrianna M. Caro, Benjamin M. Gansemer, Zhenshen Zhang, Steven H. Green

## Abstract

Spiral ganglion neurons (SGNs) constitute the sole afferent connection between cochlear hair cells and central auditory nuclei. SGNs die during postnatal developmental pruning, and also following hair cell death, which can be triggered by ototoxic agents such as aminoglycoside antibiotics, including kanamycin. After hair cell loss, animal models show extensive SGN degeneration occurring gradually over a period of weeks to months. Here, we compared spatial and temporal patterns of SGN loss and immune cell involvement in these two cases of cell death in rats. Developmental SGN pruning occurred from postnatal day 5 (P5) to P8 in the basal half of the cochlea, and from P5 to P12 in the apical half. This was accompanied by a transient increase in spiral ganglion macrophages temporally and spatially correlated with SGN death, consistent with a role clearing degenerating neurons. After deafening neonatal rats with kanamycin injections, SGN death became evident at approximately 5.5 weeks of age and persisted throughout the ganglion, with greatest loss in the middle regions; less in the base and apex. Macrophage numbers also increased but neither temporally nor spatially correlated with SGN death. Rather, increased macrophage number and activation began approximately three weeks before SGN death and was highest in the apex. Additionally, T-cells and NK cells appeared in the ganglion concurrently with SGN degeneration. These observations suggest fundamentally different roles for macrophages post-deafening than during developmental pruning and, with prior observations that anti-inflammatory drugs reduce SGN death, support a causal role for immune responses in SGN death post-deafening.

## Introduction

Sensorineural hearing loss is the most common type of hearing loss and is characterized by damage to hair cells, the sensory cells of the cochlea, or to the spiral ganglion neurons (SGNs) that conduct auditory information from the hair cells to the brain. SGNs can die from direct causes, e.g., aging, acoustic overexposure, or, secondarily, after loss of hair cells. In various animal models (e.g., rat, guinea pig, and cat), ablation of hair cells using aminoglycoside antibiotics results in significant SGN death that occurs over a period of months in rats (Alam et al., 2007; Kopelovich et al., 2013) and guinea pigs (Landry et al., 2011; Miller et al., 2007) or years in cats (Leake et al., 2011; Leake et al., 2013). The mechanisms that promote secondary SGN death after hair cell loss are currently unknown, however growing evidence implicates involvement of an inflammatory/immune response.

Macrophages are monocyte derived immune cells that function as key effector cells of the innate immune response. Macrophages are found in most tissues and organs, including the cochlea (Okano et al., 2008; Warchol, 2019). These phagocytic cells have a central role in maintaining tissue homeostasis under resting conditions and are poised to rapidly detect and respond to trauma or injury. Tissue resident macrophages are known to be present in the cochlea at rest but increase significantly in number following exposure to traumatic noise or ototoxins (Gansemer et al., 2024; Hirose et al., 2005; Kaur et al., 2015; Rahman et al., 2023). Macrophages increase in the cochlea following kanamycin-induced hair cell loss, but the timing of their increase relative to consequent SGN death remains unknown. If the increase is concomitant with, or subsequent to, the onset of neuronal death, the increased number of macrophages in the ganglion may be a response to SGN death for the purpose of phagocytizing dying neurons. Alternatively, if the increase in macrophage number in the ganglion precedes the onset of neuronal death, it is likely that macrophages have a role in SGN death other than phagocytosis. To address this, we used immunofluorescence to establish a time course of the changes in the population of immune cells—macrophages and lymphocytes—present within the spiral ganglion, from the start of kanamycin exposure and through the subsequent period of SGN death in rats.

Macrophages have also been implicated in developmental programmed cell death (PCD), a well-established mechanism for pruning of supernumerary neurons and synapses in the developing nervous system (Blaschke et al., 1996; Oppenheim, 1991). PCD is an important mechanism in the early postnatal development of the rodent cochlea, evident in remodeling of the greater epithelial ridge (Kamiya et al., 2001) and in the pruning of supernumerary SGNs and ribbon synapses (Kamiya et al., 2001; Rueda et al., 1987; Yu et al., 2021). Macrophages transiently increase throughout various regions of the cochlea during these periods of developmental remodeling and pruning (Borse et al., 2021; Dong et al., 2018; Yu et al., 2021), although, in these studies, macrophages were found not to be necessary for developmental cell death in the greater epithelial ridge (Borse et al., 2021) nor for developmental pruning of ribbon synapses (Yu et al., 2021).

To probe the role of macrophages in developmental SGN death and in SGN death post-deafening, we used immunofluorescence imaging to, first, clarify the time course of SGN death in the neonatal rat, and second, establish the time course of changes in macrophage number relative to the timing of SGN death during and following the period of normal developmental programmed neuronal death.

## Materials and Methods

### Animals and aminoglycoside deafening procedure

All animals used were Sprague-Dawley rats from our breeding colony or from pregnant Sprague-Dawley dams purchased from Envigo. The day of birth was designated as postnatal day 0 (P0). Both male and female rats were used. Rats were housed under a 12-hour light/dark cycle with ad libitum access to food and water. Protocols and procedures for all animal experiments were approved by the University of Iowa Institutional Animal Care and Use Committee.

Neonatal rats were deafened by daily intraperitoneal injections of kanamycin sulfate (400mg/kg) from postnatal day 8 (P8) to P16. As previously reported, this protocol results in the loss of outer hair cells by the end of kanamycin treatment, while inner hair cell loss is complete by approximately P20 (Bailey & Green, 2014). Deafening was verified by lack of remaining hair cells detected by immunofluorescence. Any animals with surviving hair cells were excluded from the study.

### Cochlear histology preparation

Cochleae from rats of ages P5 to P70 were used for histological analysis of hair cells, neurons, and leukocytes. Rats were deeply anesthetized with ketamine/xylazine and transcardially perfused with ice-cold PBS followed by ice-cold 4% paraformaldehyde in 0.2 M phosphate buffer. Spleens were harvested from some animals for use as positive controls for immunodetection of leukocyte markers. Cochleae were dissected from temporal bones and fixed in 4% paraformaldehyde at 4°C for up to 24 hours, followed by decalcification for 4-10 days in 0.12M EDTA, with EDTA refreshed approximately every 3 days until decalcification was complete. Cochleae were cryo-protected by sequential 30-minute incubations in 10%, 15%, 20%, and 25% sucrose, then overnight in 30% sucrose in PBS. Cochleae were then transferred to O.C.T. compound (Tissue-Tek) and maintained on a rotator overnight, then embedded in O.C.T. compound, oriented for sectioning, and flash-frozen in liquid nitrogen. Cochleae were stored at −80°C until sectioning. Serial sections (25 μm in thickness) were cut parallel to the mid-modiolar (central) plane on a Leica Cryostat and collected on Fisher Superfrost slides. Sections were stored at −20°C prior to immunolabeling.

### Immunofluorescence

A minimum of three mid-modiolar sections separated by 50 μm (i.e., every third section) were used for immunofluorescence imaging and cell counting. Immunofluorescence labeling was performed as described in Gansemer et al. (2024). Briefly, cochlear sections were permeabilized with 0.5% Triton X-100 in Tris-buffered saline pH 7.6 (TBS) for 20 minutes, then covered with blocking buffer (5% goat serum, 2% bovine serum albumin, and 0.02% sodium azide in TBS). After blocking for 3-4 hours at room temperature, sections were incubated in primary antibody in blocking buffer overnight at 4°C. After washing with TBS, sections were then incubated in secondary antibodies in blocking buffer for 3-4 hours at room temperature. Nuclei were stained with Hoechst 33342 (10 μg/ml in PBS, Sigma). Slides were coverslipped with Fluoro-Gel mounting medium in Tris-buffer, sealed with nail polish, and stored at 4°C for up to 96 hours until imaging. See Table 1 for a complete list of primary antibodies and corresponding secondary antibodies with catalog numbers and RRID’s.

**Table 1.** List of antibodies used for immunofluorescence.

| cell type | antigen | primary antibody | dilution | vendor | cat. # | secondary Ab | fluorophore | *cat. # |
| --- | --- | --- | --- | --- | --- | --- | --- | --- |
| neurons | βIII-tubulin | TuJ1, mouse IgG2a | 1/400 | Biologend | 80122 | Goat anti mouse IgG2a | Alexa Fluor 633 | A21136 |
| macrophages | Iba1 | rabbit polyclonal | 1/600 | Wako | 019-19741 | Goat anti rabbit IgG | Alexa Fluor 488 | A11008 |
| activated macrophages | CD68 | ED1, mouse IgG1 | 1/200 | Abcam | 31360 | Goat anti mouse IgG1 | Alexa Fluor 546 | A21123 |
| hair cells | myosin VI<br>myosin VIIa | rabbit polyclonal<br>rabbit polyclonal | 1/600<br>1/600 | Sigma<br>Proteus | KA-15<br>25-6790 | Goat anti rabbit IgG | Alexa Fluor 488 | A11008 |
| leukocytes | rat CD45 | OX-1, mouse IgG1 | 1/100 | Biologend | 202201 | Goat anti mouse IgG1 | Alexa Fluor 546 | A21123 |
| helper T cells | CD4 | W3/25, mouse IgG1 | 1/100 | Bio-Rad | MCA55R | Goat anti mouse IgG1 | Alexa Fluor 546 | A21123 |
| cytotoxic T cells | CD8 | OX-8, mouse IgG1 | 1/200 | Biologend | 201701 | Goat anti mouse IgG1 | Alexa Fluor 546 | A21123 |
| NK cells | CD161 | 10/78, mouse IgG1 | 1/100 | Bio-Rad | MCA1427GA | Goat anti mouse IgG1 | Alexa Fluor 546 | A21123 |
| B cells | CD19 | Clone 771404, mouse IgG2b | 1/200 | R&D | MAB7489 | Goat anti mouse IgG2b | Alexa Fluor 633 | A21146 |
| antigen presenting cells | MHCII | OX-6, mouse IgG1 | 1/100 | Biologend | 50-165-477 | Goat anti mouse IgG1 | Alexa Fluor 546 | A21123 |

\*Alexa Fluor secondary antibodies purchased from Molecular Probes

### Imaging and quantitation

Imaging and quantitation were performed as we have previously described (Gansemer et al., 2024; Rahman et al., 2023). Labeled sections were imaged on a Leica SPE confocal using a 40X (0.95 NA) or 63x (1.4 NA) objective, 1x digital zoom, and z-axis increments of 1 μm. Fiji/ImageJ (NIH) was used for all image analysis. All image stacks were assigned a random 8-digit code number using a proprietary Fiji plugin to blind the counter from the experimental condition. Cell counts were done on maximum intensity z-projections of the image stacks. Only cells with a visible nucleus were counted.

The sections of the cochlea were cut parallel to the mid-modiolar (central) plane. In each section, we assessed the spiral ganglion, which is contained within Rosenthal’s canal, in (from base to apex) the basal turn (termed “base”), the basal part of the middle turns (“mid 1”), the apical part of the middle turns (“mid 2”) and the apical turn (“apex”). The former two are collectively referenced as “basal half,” and the latter two as “apical half.” Because of the spiral anatomy of the cochlea, in sections at or close to the mid-modiolar plane at least three turns, i.e., three cross-sections, of the spiral ganglion are visible in each section (Supplementary Figure 1). The outlines of each turn were traced and the cross-sectional area of each measured. The cell count in each cross-section of Rosenthal’s canal was divided by the cross-sectional area to calculate the density.

### Statistical analysis

Statistical analyses for all data were performed using GraphPad Prism. Pairwise comparisons were made between control and KAN treated groups for each time point at each cochlear location. A Shapiro-Wilk normality test was performed for all data sets. An unpaired t-test was used for parametric data; a Mann-Whitney test for nonparametric data. A p-value <0.05 was considered statistically significant.

## Results

### Spiral ganglion neurons undergo developmental pruning during the first post-natal week

We assessed the timing of developmental SGN programmed cell death in the spiral ganglion at four locations along the base to apex axis (representative examples are shown in Figure 1). These locations, defined in Methods, are termed base and mid 1, collectively referred to as the basal half, and the mid 2 and apex, collectively referred to as the apical half. When analyzed by location, we determined that developmental SGN pruning occurs between postnatal day 5 (P5) and P8 across all regions of the spiral ganglion (Figure 2A). From P5-P8, SGN density was significantly decreased by 27.1% in the base, by 17.3% in the middle (mid) 1 region, by 21.5% in the middle (mid) 2 region, and by 20.1% in the apex. This was an overall 21.5% average decrease P5-P8 across all locations. (Rueda et al., 1987) reported an almost identical ∼22% decrease in total SGN number over the entire spiral ganglion, occurring almost entirely between P5-P6. Although not reported by Rueda et al., we found that SGN death continued past P8; the decline was significant only in the apical half of the spiral ganglion, in the basal half, there was a modest decline that was not statistically significant (Figure 2A). In this second period of SGN death from P8-P12, an additional 20.5% of SGNs were lost in the mid 2 region (p=0.008), and 14.9% were lost in the apical region (p=0.002). After P12, SGN density stabilized across all locations with no significant changes through P70. These data show that the extent and time course of SGN postnatal pruning varies by location, thus highlighting the need to attend to regional differences along the tonotopic axis.

**Figure 1.**
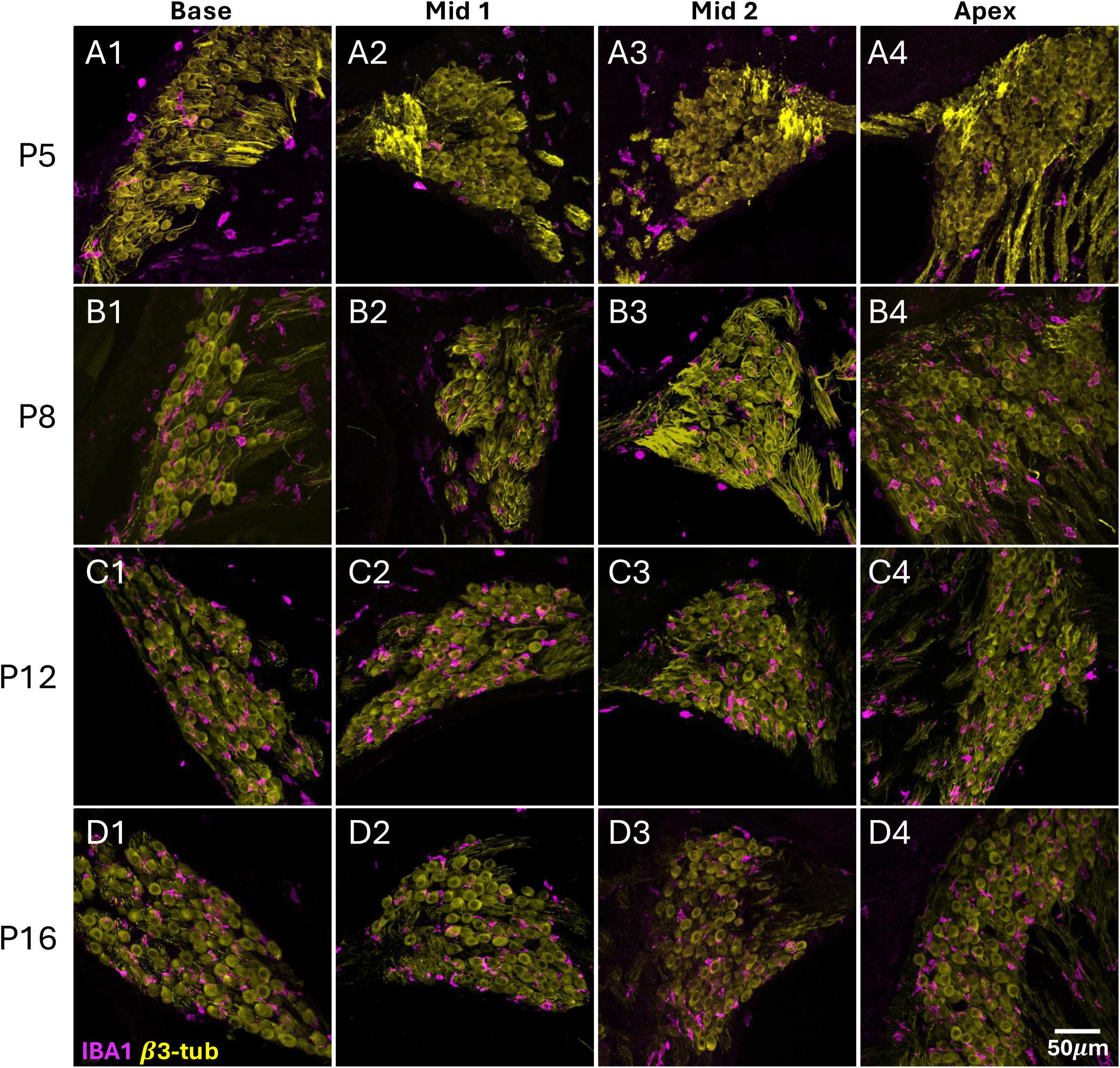
SGNs and macrophages during early postnatal developmental pruning. Shown are SGNs (β_III_-tubulin, yellow) and macrophages (IBA1, magenta) in normal control Sprague-Dawley rats at different postnatal ages, in rows (**A-D**) from top to bottom: postnatal day 5 (P5, **A1-A4**), P8 (**B1-B4**), P12 (**C1-C4**), and P16 (**D1-D4**). Representative images from cochlear cross-sections at four different locations in the spiral ganglion along the base-to-apex (tonotopic) axis are in columns from left to right: column 1 corresponds to the base (column 1), mid 1 (column 2), mid 2 (column 3), and apex (column 4).

**Figure 2.**
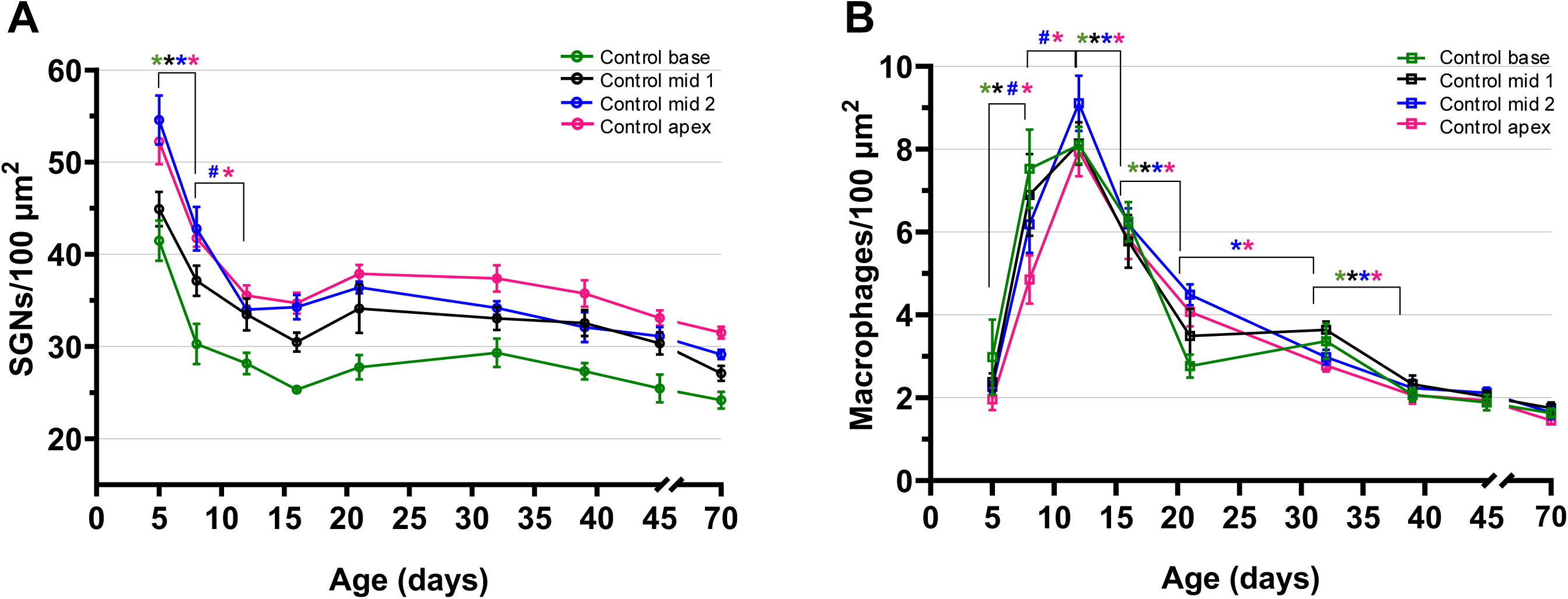
Quantitation of SGNs and macrophages in the spiral ganglion in normal control rats. Number of (**A**) SGNs (β_III_-tubulin+ cells) and (**B**) macrophages (IBA1+ cells) per 100 µm^2^ at four locations in the spiral ganglion. Shown are means ± SEM. Statistical comparisons between different timepoints are shown for each spiral ganglion location – base, mid 1, mid 2, and apex. For parametric data, significance of differences was determined using unpaired t-tests (indicated by * for p<0.05); for nonparametric data, significance of differences was determined using Mann-Whitney tests (indicated by # for p<0.05). The symbols for significance are color-matched to the corresponding cochlear locations, as indicated in the legend. For all timepoints and conditions, n=5-8 animals.

### Macrophages in the spiral ganglion transiently increase in number during the first two post-natal weeks

Concomitant with neonatal SGN pruning, spiral ganglion macrophage number increased in neonatal rats. We observed a significant increase in macrophage density from P5 to P8 throughout the spiral ganglion, varying by location (Figure 2B). There was a 151.8% increase in the base, 189.2% increase in the mid 1 region, 173.1% increase in the mid 2 region, and 147.5% increase in the apex. From P8 to P12, there was an additional 47.2% increase in macrophage density in the mid 2 region (p=0.03) and 63.5% increase in the apex (p=0.003). The greater increase in the apex relative to the base may be related to the more extended period of developmental SGN death in the apical half relative to the basal half (Figure 2A). In the developing cochlea, macrophage number peaked at P12 across all regions and gradually decreased with age through P39, after which macrophage density remained nearly constant at approximately 2 cells/100 µm^2^ (Figure 2B). Macrophages are known to be involved in the clearance and phagocytosis of dying cells. That their increase in number closely correlates temporally and spatially with developmental SGN pruning, is consistent with this function. Following the completion of SGN pruning, the remaining macrophages constitute a stable resident population in the normal spiral ganglion, absent any trauma to the cochlea.

### Neurons die throughout the spiral ganglion after kanamycin-induced hair cell loss

In the rat, daily systemic injection of the aminoglycoside antibiotic kanamycin sulfate from P8-P16 results in complete loss of inner and outer hair cells, from P14 to P21 (Bailey & Green, 2014). Following hair cell loss is a gradual degeneration of SGNs over the course of approximately 12 weeks, ultimately resulting in the death of over 90% of SGNs relative to age-matched controls (Alam et al., 2007). Examples are shown in Figure 3. However, SGN degeneration is not immediate after hair cell loss. Here, we found that the onset of significant SGN death in kanamycin-treated (KAN) rats in the basal (Figure 4A) and apical (Figure 4B) regions of the ganglion was between P32 and P39, two to three weeks after death of all hair cells. By P39, SGN density was significantly decreased in KAN rats compared to controls by 18.5% in the mid 1 region, by 16.6% in mid 2, and by 19.3% in the apex. In the base, SGN density was decreased by 9.2% in KAN rats and was not significantly different from control rats (p=0.09). By P45, SGN loss was significant in all regions: relative to age-matched control hearing rats, SGN density was decreased by 15.2% in the base, by 17% in mid 1, by 19.6% in mid 2, and by 15.6% in the apex. By P70, SGN density in KAN rats was significantly decreased, relative to age-matched control hearing rats, by 22.7% in the base, by 46.8% in mid 1, by 44% in mid 2, and by 32.1% in the apex compared to age-matched controls. Thus, the middle regions of the cochlea had the greatest SGN loss, with more modest SGN loss at the base and apex. Thus, as was the case for developmental neuronal death, the rate of neuron loss varied spatially in the ganglion along the base-to-apex axis, although the spatial patterns differ.

**Figure 3.**
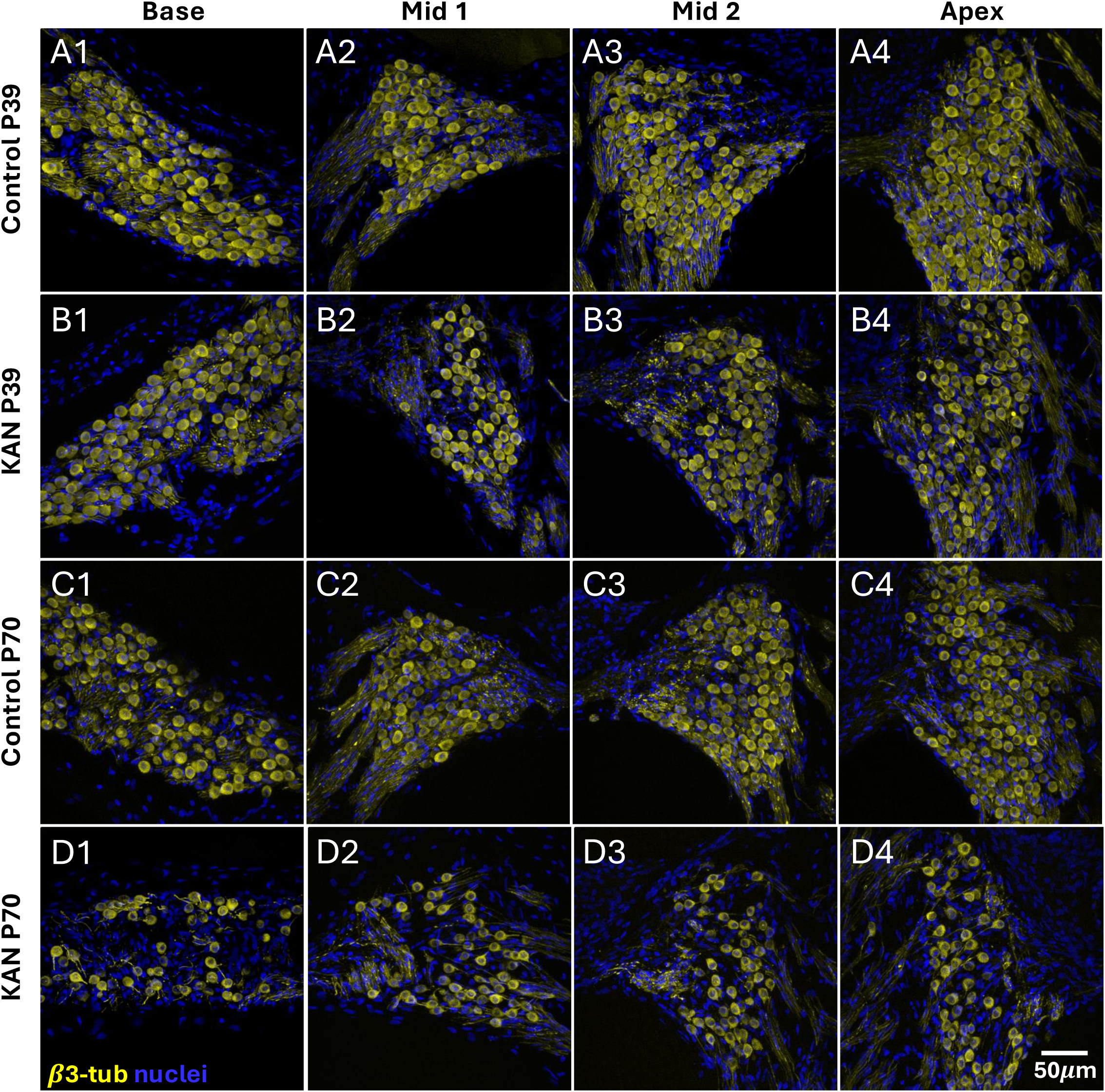
SGNs in adult control and kanamycin-treated rats. (A-D) Representative images showing SGNs (β_III_ -tubulin, yellow) and all nuclei (Hoechst 33342, blue) at P39, when significant SGN death is first detectable post-deafening, in hearing control **(A1-A4)** and kanamycin treated (KAN, **B1-B4**) rats. Representative images at P70 in control **(C1-C4)** and KAN treated **(D1-D4)** rats.

**Figure 4.**
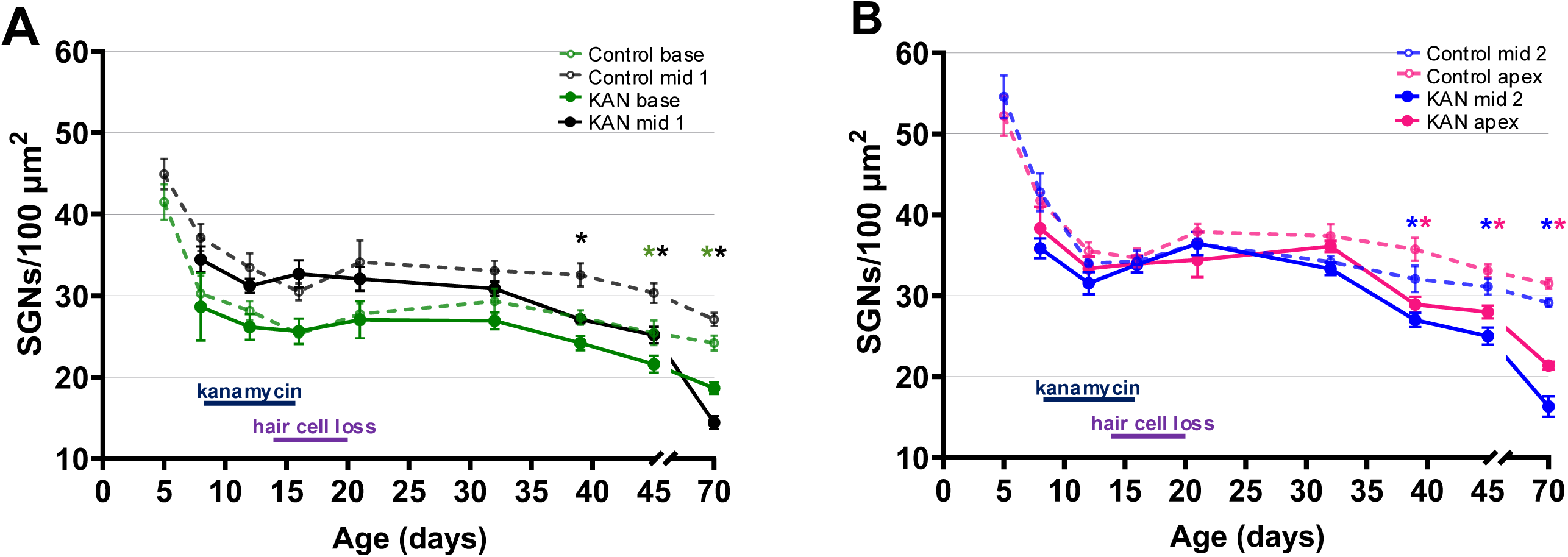
Quantitation of SGNs by cochlear location in adult control and kanamycin-treated rats. Shown are means ± SEM for measures of SGN density (SGNs/100 µm^2^) in the **(A)** basal half (base and mid 1) and **(B)** apical half (mid 2 and apex) of KAN and control hearing rats at various timepoints over the period P5-P70. Data from control rats are from Figure 2, here replotted to be compared directly to KAN treated rats. Comparisons were made between age-matched KAN and control rats at each location with significance determined by unpaired t-tests, indicated by * for p<0.05; color-coded to location. n=4-10 animals for all timepoints and conditions. Colored bars on the timeline show the timing of kanamycin treatment (dark blue, P8-P16) and resulting hair cell loss (purple, approx. P12-P19 (Bailey & Green, 2014). A significant decline in SGN number in KAN rats, relative to control rats, is first detectable at P39.

### Macrophages in the spiral ganglion increase in number and activation prior to onset of post-deafening SGN loss

As shown in Figure 2, macrophage density in control rats transiently increased within the spiral ganglion concomitantly with normal developmental pruning. In contrast, in KAN rats, we found that macrophage number increased prior to the onset of post-deafening SGN death and persisted for several weeks (Figures 5, 6). However, the time course of changes in macrophage density varied spatially and quantitatively within the ganglion, particularly between the basal and apical halves of the ganglion. In the basal half, spiral ganglion macrophage number in KAN-treated rats followed the normal developmental temporal pattern through P16 (Figure 6A). A significant difference between KAN and control rats was not apparent in the basal half until P21, when there was a significant increase in macrophage abundance in KAN rats that persisted through P70. In the basal half, macrophage density in KAN rats gradually declined from P32 to P70, from ∼5 cells/100 µm^2^to ∼3 cells/100 µm^2^, while remaining significantly increased relative to controls.

**Figure 5.**
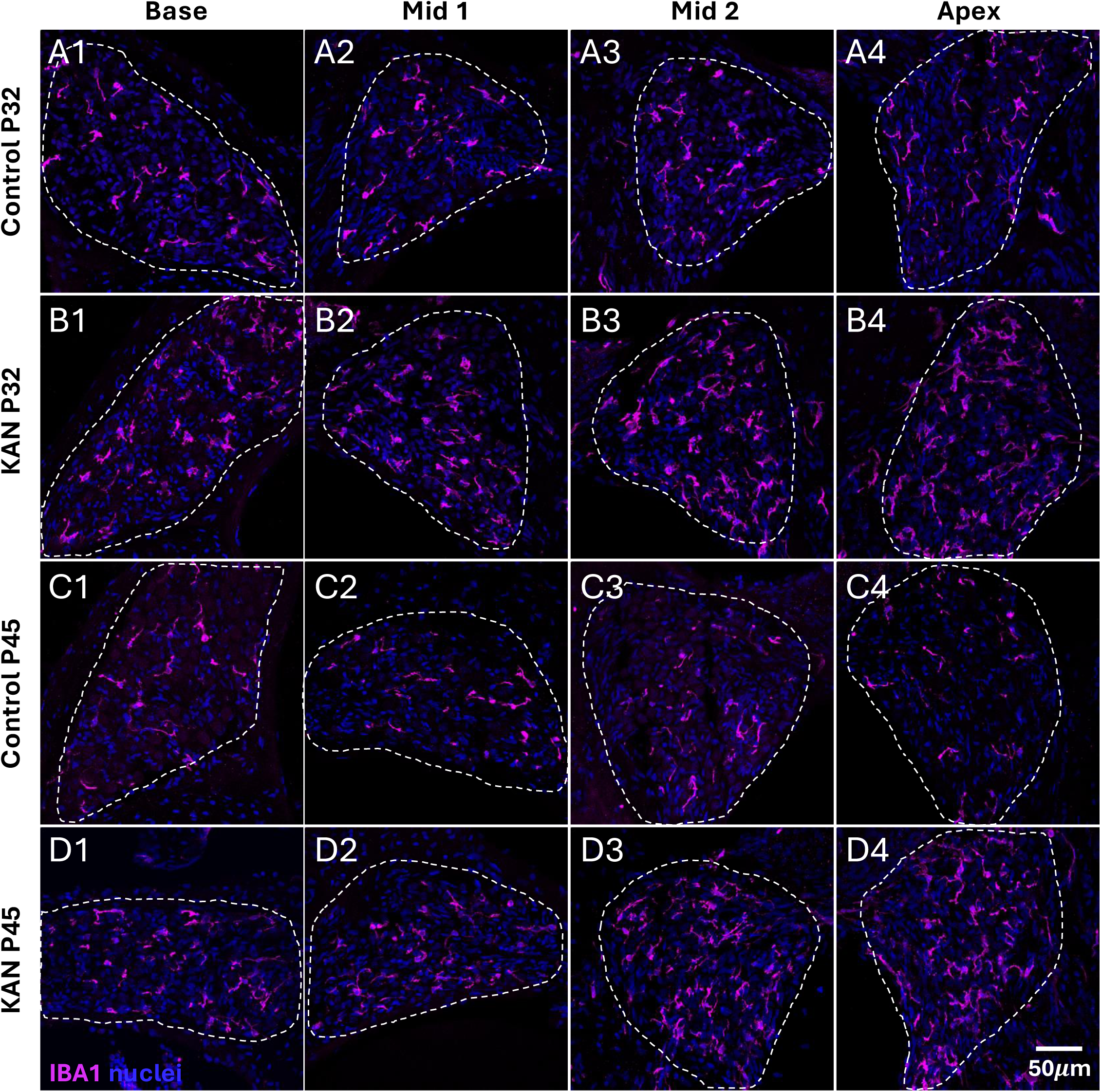
Macrophages in adult control and kanamycin-treated rats. Representative images of macrophages (IBA1, magenta) and all nuclei (blue) from different cochlear locations, left to right: base (column 1), mid 1 (column 2), mid 2 (column 3) and apex (column 4). Dotted outlines delineate Rosenthal’s canal. **A**,**B:** Images from hearing control **(A1-A4)** and kanamycin treated (KAN, **B1-B4)** rats at **P32**, prior to onset of SGN death. **C**,**D:** Images from hearing control **(C1-C4)** and kanamycin treated (**D1-D4)** rats at **P45**, when SGN death is underway.

**Figure 6.**
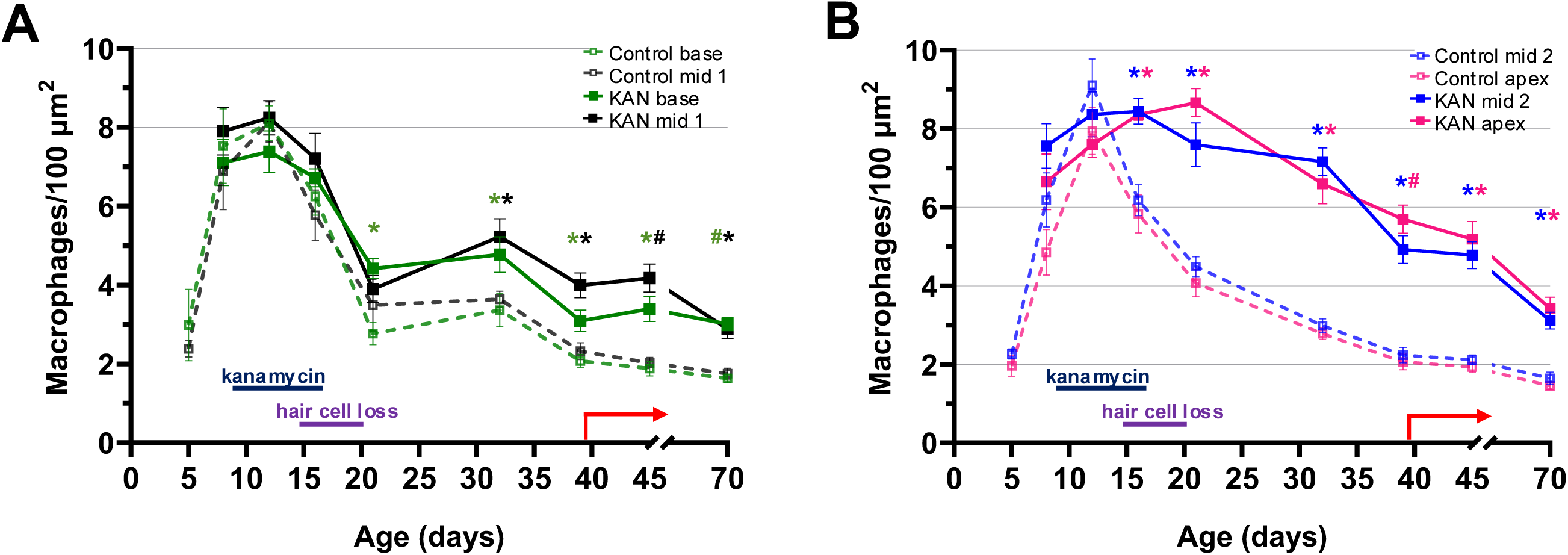
Quantitation of macrophages by cochlear location in adult control and kanamycin-treated rats. Shown are means ± SEM for measures of macrophage density (macrophages/100 µm^2^) in the **(A)** basal half (base and mid 1) and **(B)** apical half (mid 2 and apex) of KAN and control hearing rats at various timepoints from postnatal days 5-70. Data from control rats are from Figure 2, here replotted to be compared directly to KAN treated rats. Comparisons were made between age-matched KAN and control rats at each location. For parametric data, significance of differences was determined using unpaired t-tests (indicated by * for p<0.05); for nonparametric data, significance of differences was determined using Mann-Whitney tests (indicated by # for p<0.05). The symbols are color-matched to location. n=4-10 animals for all timepoints and conditions. As in Figure 4, colored bars on the timeline show the timing of kanamycin treatment and resulting hair cell loss and a red arrow indicates the onset of SGN death. Macrophage number increases approximately at the time of hair cell death and precedes the onset of SGN death.

In contrast, in the apical half of the cochlea, what was observed in KAN rats was maintenance of the increase in macrophages initiated during developmental SGN pruning (Figure 6B). That is, macrophage number did not decrease following the developmental peak at P12, as in control rats. Rather, the increase was maintained through P21, after which macrophage number gradually declined, but remained significantly elevated relative to controls and to the basal half in KAN rats through P70. By P70, the decline in macrophage number in the apical half had accelerated, bringing the density in the basal and apical halves close to equality, with approximately ∼3 cells/100 µm^2^ at all cochlear locations.

We then asked whether the increase in the number of macrophages is associated with an increase in macrophage activation. We used CD68, a lysosomal marker indicative of phagocytic activity, as an indicator of macrophage activation (Chistiakov et al., 2017; Holness et al., 1993). We have previously shown an increase in CD68 gene expression and CD68 immunoreactivity in macrophages by P70 in the spiral ganglia of KAN rats (Gansemer et al., 2024; Rahman et al., 2023). Here, we show that in KAN rats, this increase in CD68 immunoreactivity in spiral ganglion macrophages (examples of which are shown in Figures 7A-C) begins as early as P21 in the mid 1 (p=0.02) and mid 2 (p=0.02) regions of the cochlea (Figure 7D). By P32, there was a significant increase in the apex, with 18-35% of IBA1+ cells in the ganglion co-expressing CD68. This fraction increased concomitantly with the onset of significant SGN death by P39, when 43-58% of macrophages were CD68+ throughout all regions of the spiral ganglion. This increase in the proportion of activated macrophages was sustained at least through P45. This contrasts with observations in control rats, in which few spiral ganglion macrophages expressed CD68 (Figure 7D). These data indicate that macrophage activation and macrophage number increase in parallel in the spiral ganglion, starting well prior to the onset of significant SGN death. As neuronal death is not yet evident when macrophages increase in both number and activation, we infer that macrophages are not simply being recruited and activated for the purpose of phagocytizing dead/dying SGNs.

**Figure 7.**
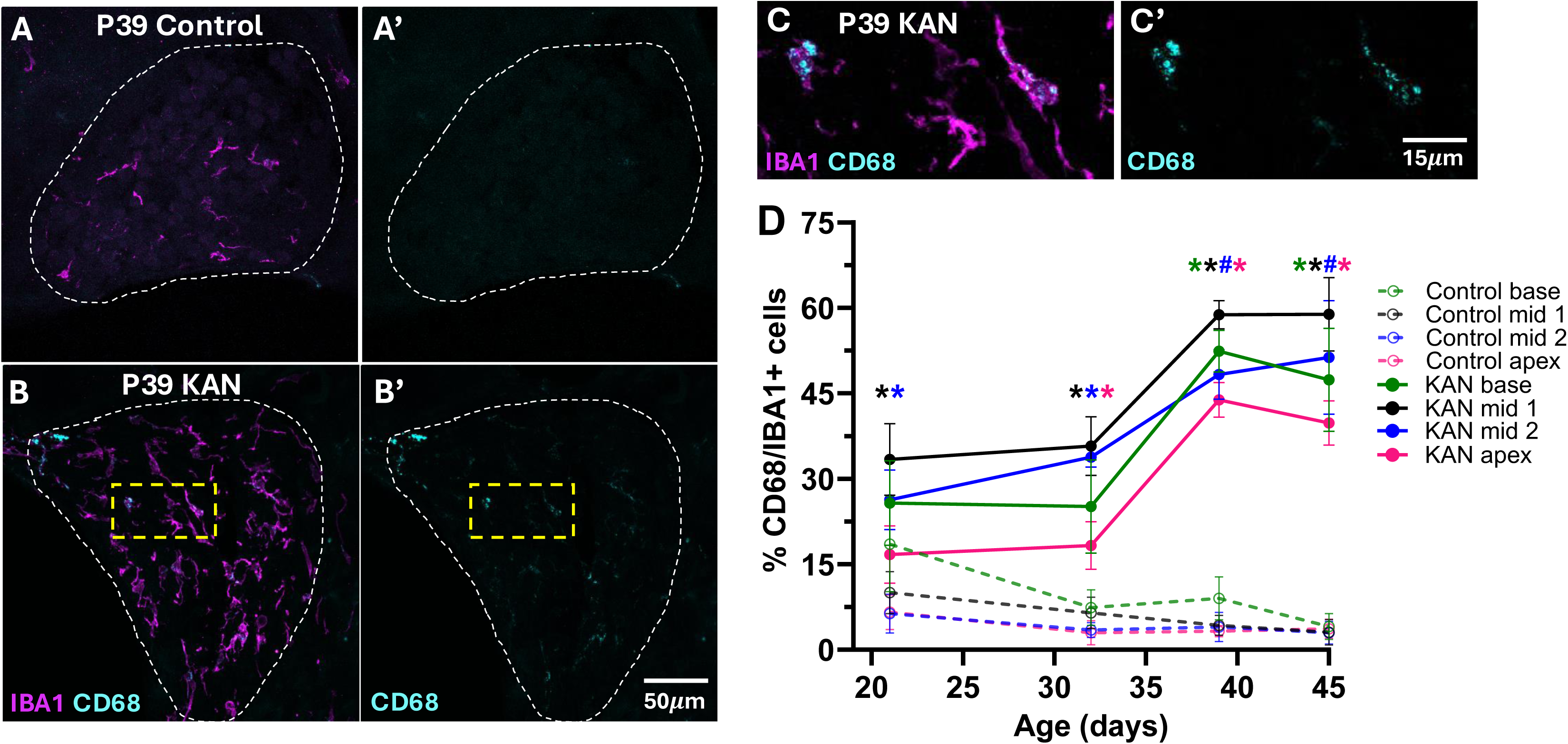
CD68 expression in spiral ganglion macrophages. Representative images from the mid 2 region of spiral ganglia from a P39 control rat (**A**, **A’**) and a kanamycin-treated rat (**B**, **B’**, **C**, **C’**). C and C’ are higher magnification images. The dashed yellow square outline in **B** outlines the area shown at higher magnification in **C.** These images show labeling with IBA1 (magenta) and CD68 (cyan), with the two labels superimposed in **A**, **B**, and **C**, but CD68 only in **A’**, **B’**, and **C’**. Dotted outlines delineate Rosenthal’s canal in **A**, **A’**, **B**, and **B’**. The percentage of IBA1-positive cells that are also CD68-positive (double positive cells) is shown in **D**. Shown are means ± SEM. Comparisons were made between age-matched KAN and control rats at each location with significance for parametric data determined by unpaired t-tests, indicated by * for p<0.05, or, for nonparametric data, by Mann-Whitney tests, indicated by # for p<0.05. The symbols are color-matched to location. n=4-6 animals for all timepoints and conditions. The increase in macrophage number is accompanied by a significant increase in the fraction of macrophages that are activated.

### MHCII expression increases in spiral ganglion macrophages post-deafening

To explore the function of macrophages in the spiral ganglion post-deafening, we assessed MHCII expression in the macrophages. Examples are shown in Figures 8A,B. We found that in KAN rats, at P21, the total number of MHCII+ macrophages (IBA1+/ MHCII+) declined in the basal half (Figure 8C) of the ganglion (base, ns; mid 1, p=0.007) without significant change in the apical half at P21. However, by P32, in the apical half of the ganglion the total number of MHCII+ macrophages in KAN rats had significantly increased relative to control hearing rats (Figure 8C’) and remains elevated thereafter. In contrast, in the basal half of the ganglion, the number of macrophages expressing MHCII in KAN rats is not significantly elevated relative to control rats at any time post-deafening.

**Figure 8.**
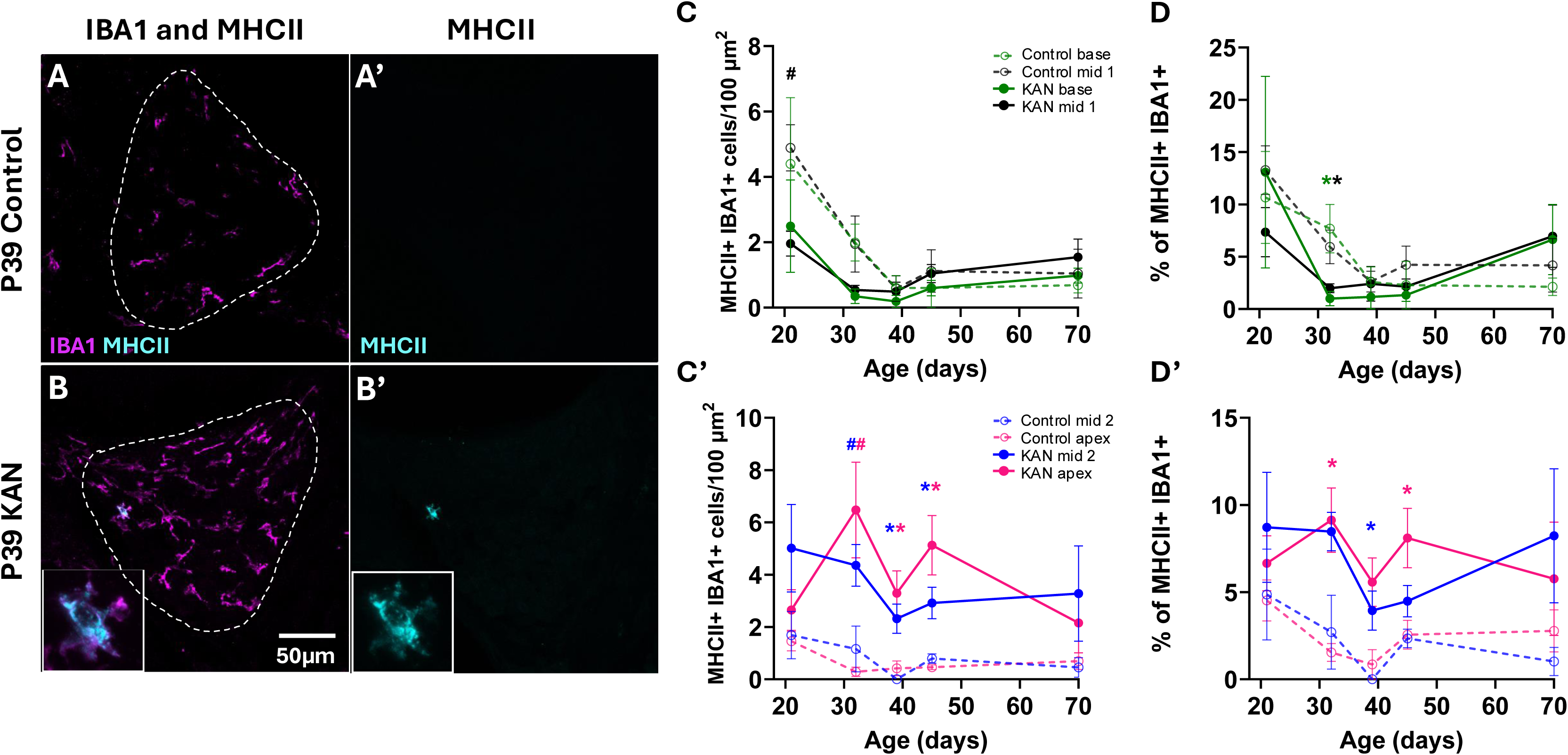
MHCII expression in spiral ganglion macrophages. Representative images from the mid 2 region of spiral ganglia from a P39 control rat (**A**, **A’**) and a kanamycin-treated rat (**B**, **B’**). These images show labeling with IBA1 (magenta) and MHCII (cyan), with the two labels superimposed in **A** and **B**, but MHCII only in **A’** and **B’**. Dotted outlines delineate Rosenthal’s canal in **A** and **B**. Higher magnification images are shown in the insets in **B** and **B’**. The number (means ± SEM) of MHCII-positive macrophages is plotted in **C** and **C’**. The percentage (means ± SEM) of macrophages that are also MHCII-positive (double positive cells) is plotted in **D** and **D’**. **C** and **D** show basal half (base and mid 1); C’ and D’ show apical half (apex and mid 2). Comparisons were made between age-matched KAN and control rats at each location with significance for parametric data determined by unpaired t-tests, indicated by * for p<0.05, and, for nonparametric data, by Mann-Whitney tests, indicated by # for p<0.05. The symbols are color-matched to location. n=4-6 animals for all timepoints and conditions. The increase in macrophage number is accompanied in the apical half by a significant increase in the fraction of macrophages that express MHCII.

While the increase in number of MHCII+ macrophages can be explained, at least in part, by the increase in the total number of macrophages, we further found that the *fraction* of macrophages expressing MHCII also increased significantly in the apex by P32 (Figure 8D, D’). This increase was already apparent prior to the start of significant SGN death at P39 and persisted through P45 (Figure 8D’), i.e., through the early period of SGN death. This increased MHCII immunofluorescence is consistent with our previously reported microarray and RNAseq studies (Gansemer et al., 2024; Rahman et al., 2023) that showed increased gene expression of MHCII and MHCII-related components in the spiral ganglia of kanamycin-deafened rats. While significantly elevated compared to age-matched control hearing rats, the total number of MHCII+/IBA1+ cells – 0.22-0.65 cells/100 µm^2^ – in the apical half of the spiral ganglion of KAN rats is still <10% of all macrophages. Nevertheless, this does indicate acquisition of novel functions by spiral ganglion macrophages post-deafening. Because MHCII is associated with antigen presentation to and activation of CD4+ helper T cells, we asked whether lymphocytes, including T cells, are present within the ganglion, and if so, when and where they appear relative to the period of post-deafening SGN loss.

### Non-macrophage immune cells infiltrate the ganglion post-deafening

While macrophages are the most abundant immune cell type responding to damage in the cochlea, other leukocytes have been shown to increase in number following damage in both mouse and rat models (Gansemer et al., 2024; Hirose et al., 2005; Rai et al., 2020), although in smaller numbers than the macrophages. Here we asked about the timing of appearance in the spiral ganglion of immune cells other than macrophages. To identify such cells, we used the pan-leukocyte marker CD45 in combination with the macrophage marker IBA1. CD45+/IBA1– cells were classified as non-macrophage immune cells, examples of which are shown in Figures 9A-C. Such CD45+/IBA1– cells were rarely seen in spiral ganglia from control rats (Figure 9A,G), indicating that, normally, nearly all resident immune cells in the spiral ganglion are macrophages. In KAN rats, we found a significant increase in the number of CD45+/IBA1-cells in the mid 1, mid 2, and apex regions of the spiral ganglion starting by P39 (Figure 9 B-C,G), which is concomitant with the onset of SGN death. The total number of these cells was relatively small, at a density of 0.21-0.40 cells/100 µm^2^ in P39 KAN rats, varying with location – relative to 3.1-5.7 cells/100 µm^2^ for macrophages and 24-27 cells/100 µm^2^ for SGNs – but was significantly increased relative to controls. Thus, the number of non-macrophage immune cells is between 4% and 13% of the number of macrophages, consistent with the report of Hirose et al. (2005) that ∼5% of the immune cells recruited to the cochlea following acoustic trauma are not CX3CR1+ macrophages. We found that this population of non-macrophage immune cells remained significantly elevated in density in all regions of the spiral ganglion through P70 in KAN rats.

**Figure 9.**
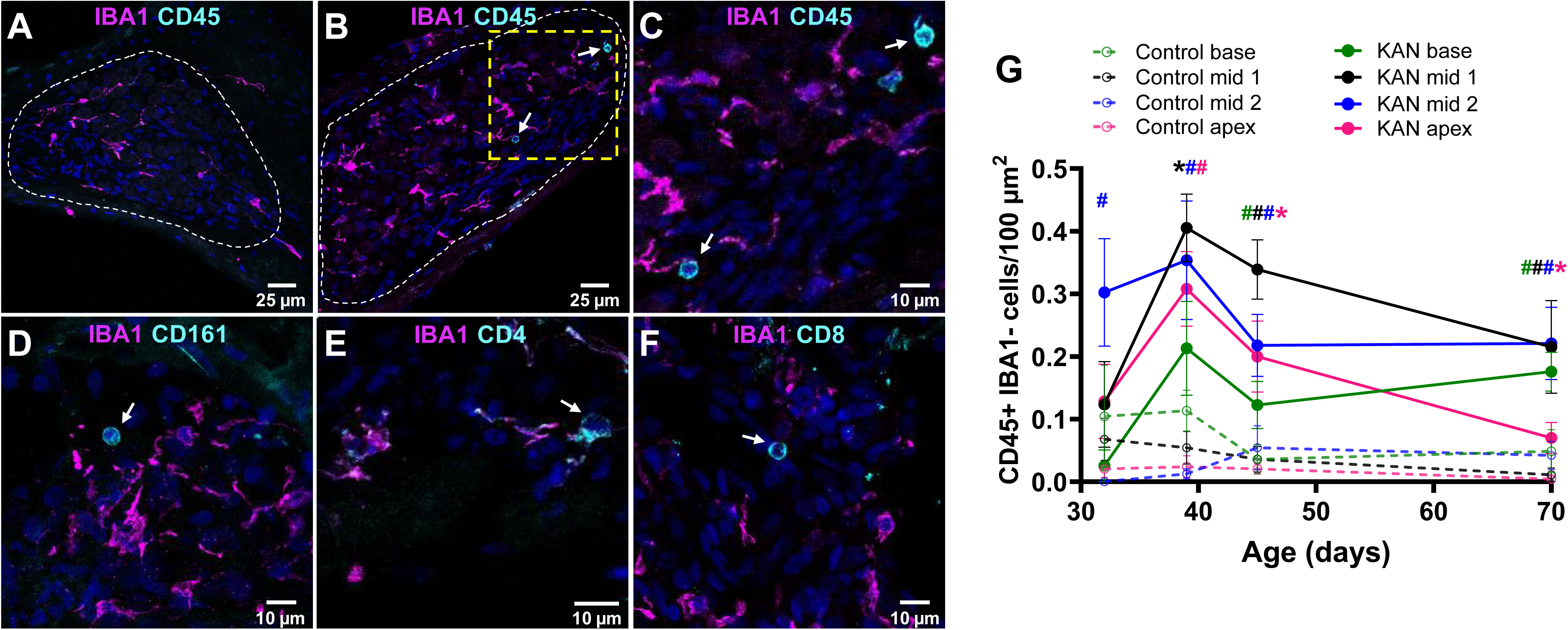
A heterogenous population of non-macrophage immune cells increases in the spiral ganglion after hair cell loss. **A**,**B**,**C:** Representative spiral ganglion images from the Mid 1 region of a P39 control rat (**A**) and a P39 kanamycin-treated (KAN) rat (**B**,**C**) labeled to show presence of CD45+/IBA1– leukocytes in the ganglion of the KAN rat, but not in the control rat. Images in **C**-**F** are at higher magnification. In A and B, a dotted outline delineates Rosenthal’s canal. The dashed yellow square in **B** outlines the area shown at higher magnification in **C**. The population of CD45+/IBA1– lymphocytes in KAN rats includes CD161A+ NK cells (**D**), CD4+ helper T cells (**E**), and CD8+ cytotoxic T cells (**F**). In all images, arrows mark the referenced immunolabeled lymphocytes. **G:** Number of lymphocytes (CD45+/IBA1– cells) per 100 µm^2^ at four locations in the spiral ganglion. Shown are means ± SEM. Statistical comparisons between different timepoints are shown for each spiral ganglion location – base, mid 1, mid 2, and apex. Comparisons were made between age-matched KAN and control rats at each location with significance determined, for parametric data, by unpaired t-tests, indicated by * for p<0.05, or by Mann-Whitney tests, for nonparametric data, indicated by # for p<0.05. The symbols are color-matched to location. n=6-10 animals for all timepoints and conditions.

As shown in Figure 9C, these CD45+/IBA1-cells are small and round in morphology, with large nuclei, a characteristic of lymphocyte morphology. Subsets of these cells were positive for CD4, CD8, or CD161 (Figure 9D-F), identifying, respectively, helper T-cells, cytotoxic T cells, and NK cells. We did not find cells immunoreactive with CD19, a marker for B cells (data not shown). These results indicate that both macrophages and a mixed population of lymphocytes respond to damage in the cochlea, consistent with previous reports (Gansemer et al., 2024; Hirose et al., 2014; Rai et al., 2020). However, the timing of appearance of these different types of immune cells is substantially different. Macrophages appear in the spiral ganglion and are activated while hair cells are dying, approximately three weeks prior to the onset of SGN death and persist in elevated numbers during the period of SGN death. In contrast, lymphocytes appear only later, at the time that SGNs are starting to die in significant numbers. Furthermore, the presence of these lymphocytes in the ganglion raises the possibility that MHCII-expressing macrophages may be participating in antigen presentation to T cells.

## Discussion

Here we used immunofluorescence to establish a time course of the changes in the population of immune cells present within the spiral ganglion during two periods of SGN death: first, during developmental pruning and second, associated with kanamycin-induced hair cell loss. These two instances of neuronal death show distinctly different spatiotemporal patterns of immune system involvement. There is a transient increase in macrophage number during and immediately following the period of postnatal pruning across all regions of the ganglion. The macrophages that remain are resident cells that display a resting phenotype (IBA1+/CD68-) (Figure 7) and are the primary immune cell type present in the spiral ganglion of normal hearing rats. In the spiral ganglion of KAN rats, we found a significant increase in the number of macrophages relative to controls as early as P16 in the apical half and by P21 in the basal half— overlapping in the apex with the increase associated with developmental pruning. As early as P21, there was a significantly larger proportion of CD68+ macrophages, indicating that there is an increase in both macrophage number and activation prior to the start of significant SGN death. Additionally, in KAN rats, we identified an increase in lymphocytes, including both CD4+ helper and CD8+ cytotoxic T cells, as well as CD161+ NK cells across the spiral ganglion that occurs concurrent with the onset of significant SGN death at P39. Thus, there is a heterogenous population of both innate and adaptive immune cells that respond to the spiral ganglion after hair cell loss, with distinct temporal and regional characteristics.

Aside from the observations of immune involvement in SGN death post-deafening, the spatial and temporal patterns of SGN death observed here in the rat are unexpected. Previous studies of SGN death post-deafening in guinea pigs and cats showed neuronal loss earliest in the basal turn, then, and gradually progressing, apically. Guinea pigs show a rapid onset of SGN loss, within hours, during the first two weeks post-deafening, and further loss during the subsequent six weeks at a slower rate (Dodson, 1997). In cats, SGN loss is likewise rapid initially with ∼60% of SGNs lost in the first few weeks post-deafening followed by a slower progressive loss, taking months to years, that is slowest in mid and apical regions with the basal turn showing the most rapid and extensive SGN loss (Leake et al., 2013). In the rat, the temporal pattern is distinctly different: we find no detectable neuronal loss until about three weeks after hair cell loss, and there is no apparent initially faster rate of loss, as shown in Figure 4. With regard to the spatial pattern, this too differs in rats from what has been reported in cats and guinea pigs. In rats, SGN loss appears to be slowest in the base and most rapid and extensive in the middle regions, relative to the base and apex (Figure 4). The rats in our study started receiving kanamycin prior to hearing onset while the guinea pigs in the Dodson (1997) study received aminoglycosides as young adults, well after hearing onset. However, the cats in the Leake et al. (2013) study also started receiving kanamycin prior to hearing onset, so it’s not clear that the timing of deafening in relation to hearing onset can explain the differences in the spatial and temporal patterns of neuronal loss among these animal models.

SGN death during postnatal pruning also varies quantitatively along the base-apex axis. Between P5 and P12, SGN density declines by 28.8% in the basal half (base and mid 1) and by 34.8% in the apical half (mid 2 and apex) (Figure 2). The close temporal correlation between SGN death and increase in macrophage number between P5 and P12 appears to be accompanied by a spatial correlation. The greater neuronal loss in the apical half is associated with an increased number of macrophages: ∼300% increase from P5 to P12 in the apical half and ∼200% increase in the basal half. These correlations are consistent with a hypothesis that the increase in macrophage number is for the purpose of phagocytosis of dying SGNs during this period of neurodevelopment.

This temporal pattern of SGN death and macrophage recruitment during postnatal pruning is distinctly different from that seen post-deafening. SGNs in KAN rats do not start to die in significant numbers until P39 (Figure 4), approximately three weeks after loss of all hair cells (Bailey & Green, 2014), while macrophage number (Figure 6) and activation (Figure 7) are already significantly increased by P21. These data are not consistent with a hypothesis that the increase in macrophage number is solely for the purpose of phagocytosis of dying neurons, as SGNs have not yet started to die in large numbers when macrophage number increases. One possibility is that the primary cue for macrophage recruitment is hair cell degeneration. Macrophages have been shown to invade the organ of Corti in response to hair cell damage from acoustic and/or ototoxic trauma (Fredelius & Rask-Andersen, 1990; Kaur et al., 2015; Manickam et al., 2023). Similarly, it is likely that macrophages here are recruited to the organ of Corti in response to kanamycin-induced hair cell death. While hair cell death can account the for recruitment of macrophages to the organ of Corti, it’s not yet clear what recruits them to the spiral ganglion prior to neuronal death.

As noted above, there is a spatial correlation of SGN death and macrophage number within the ganglion during postnatal pruning. In contrast, there does not appear to be a consistent spatial correlation between SGN death and macrophage number post-deafening. The data presented in Figures 4 and 6 are replotted in Supplementary Figure 2 to better demonstrate quantitative differences in the decrease in SGN density and increase in macrophage density by cochlear location. Macrophage number increases earliest in the apical half, which does correspond to an initially faster loss of SGNs in the apex, although the macrophage increase precedes SGN loss. Throughout most of the period of SGN death from P39 through P70, macrophage number remains highest in the apical half, although the density of macrophages does even out throughout the ganglion by P70 (Supplementary Figure 2). This does not correlate with SGN loss, which remains greatest in the middle regions of the spiral ganglion. Moreover, the fraction of macrophages expressing MHCII is higher in the apical than in the basal half of the ganglion. One possible explanation for the greater number of macrophages in the apical half of the ganglion throughout the early period of SGN degeneration is correlation with cytokine expression. Some cytokines, implicated in recruiting macrophages, and upregulated post-deafening (Gansemer et al., 2024; Rahman et al., 2023), show an initially higher level of expression in the apical half of the spiral ganglion (Rahman et al., 2023) relative to the basal half (Gansemer et al., 2024; Rahman et al., 2023). Specifically, in the entire ganglion, the CCR2 ligands CCL2 and CCL7 are upregulated, respectively, by 1.2x and 1.9x at P32 post-deafening, and by 5.1x and 3.9x at P60 (Gansemer et al., 2024; Rahman et al., 2023).

Once recruited to the spiral ganglion, the macrophages appear to be playing a causal role in SGN death. This is evidenced by studies showing that elimination of macrophages (Shimada et al., 2023) or treatment with anti-inflammatory agents that reduce macrophage activation (Rahman et al., 2023) promote SGN survival after hair cell loss. This hypothesis is consistent with the observation that macrophage recruitment and activation precede SGN death and remain elevated while SGN numbers decline. Kaur et al. (2015) has reported that SGN death post-deafening is increased by deletion of the fractalkine receptor (CX3CR1) in macrophages. This can be explained by an increase in macrophage reactivity and pro-inflammatory activity due to loss of response to the CX3CR1 ligand, the homeostasis-promoting factor CX3CL1 (Cardona et al., 2006; Iemmolo et al., 2023).

It is plausible that macrophages promote a neurotoxic environment in the spiral ganglion that contributes to neuronal death, as has been seen in examples of chronic inflammation and neurodegeneration in the central nervous system (Comitre-Mariano et al., 2024; Eitas et al., 2014; Muller et al., 2025). Animal models of traumatic brain injury have shown that macrophage and microglial activation can result in the release of reactive oxygen species, cytokines, and other proinflammatory factors that contribute to chronic neurodegeneration (Kumar et al., 2013; Loane & Kumar, 2016). Similarly, in the cochlea, macrophages may directly compromise SGN survival via comparable cytotoxic effects (Gansemer et al., 2024; Rahman et al., 2023).

Alternatively, though not mutually exclusive, macrophages may indirectly affect SGN survival through the recruitment and activation of the adaptive immune response. MHCII is a key signaling molecule used to bridge communication between cells of the innate and adaptive immune systems (Unanue, 1984). MHCII-mediated antigen presentation by phagocytes, including macrophages, activates CD4+ helper T cells and initiates adaptive immune responses (Roche & Furuta, 2015). Upregulation of MHCII (Subbarayan et al., 2020) and T cell infiltration into the brain has been associated with neuron loss in CNS neurodegenerative conditions such as Parkinson’s (Brochard et al., 2009; Subbarayan et al., 2020) and Alzheimer’s diseases (Zeng et al., 2024). The increase in the proportion of spiral ganglion macrophages expressing CD68 (Figure 7) and MHCII (Figure 8) indicates a functional shift to a state of activation. This, coupled with the presence of lymphocytes, namely CD4+ helper T cells, into the ganglion at P39 (Figure 9) raises the possibility that macrophages may activate adaptive immune responses after hair cell damage via MHCII-mediated antigen presentation. In the cochlea of noise-exposed mice, there is also an increase in both MHCII+ cells and CD4+ cells, indicative of antigen-presenting function (Yang et al., 2015). The appearance of lymphocytes concomitant with the onset of significant SGN death is consistent with the hypothesis that these cells may be contributing to SGN death post-deafening. Further studies should focus on the roles of the innate and adaptive immune systems in SGN loss after deafening.

## Acknowledgments

We thank Catherine Kane for maintaining our animal colony and Tristan Brown for assistance in labeling of cochlear sections. This work was supported by the National Institute on Deafness and other Communication Disorders grants NRSA DC021590 (AMC) and R01 DC015790 (SHG).

## Author contributions

AMC: Data curation, Formal analysis, Writing—original draft, review and editing, Conceptualization. BMG: Data collection, Conceptualization. ZZ: Data collection. SHG: Formal analysis, Writing—review and editing, Conceptualization, Project administration.

## Data availability statement

The data supporting the findings of this study are available upon reasonable request.

---

**Supplementary figure 1.**
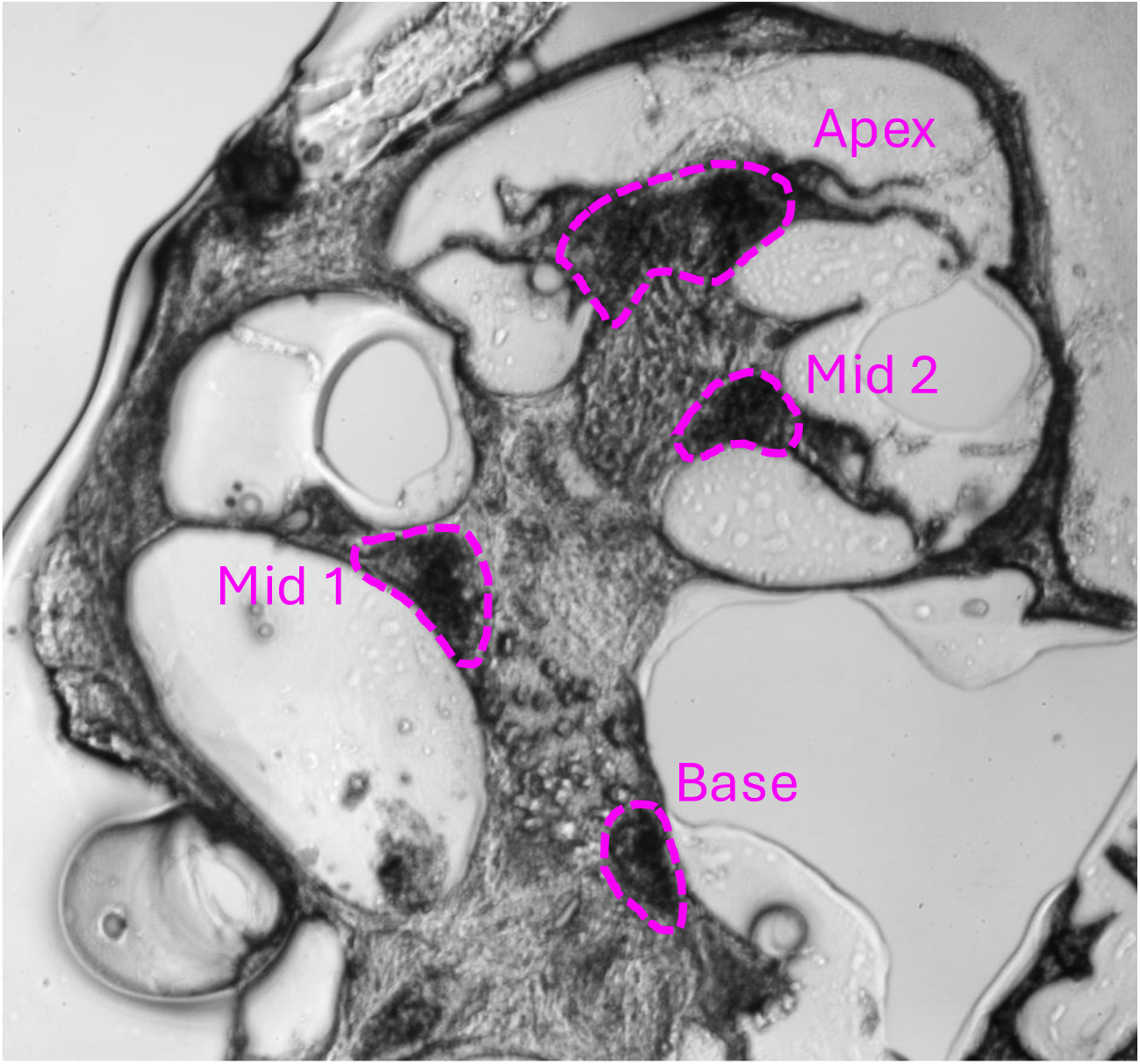
Low magnification image of a representative section through the mid-modiolar (central) plane showing four cross-sections of Rosenthal’s canal, which contains the spiral ganglion. Because of the spiral topology of the cochlea, each cross-secion is at a different region along the tonotopic (base to apex) axis. The outlines of each cross-section are outlined and labeled: base, middle (mid) 1, mid 2, and apex.

**Supplementary figure 2.**
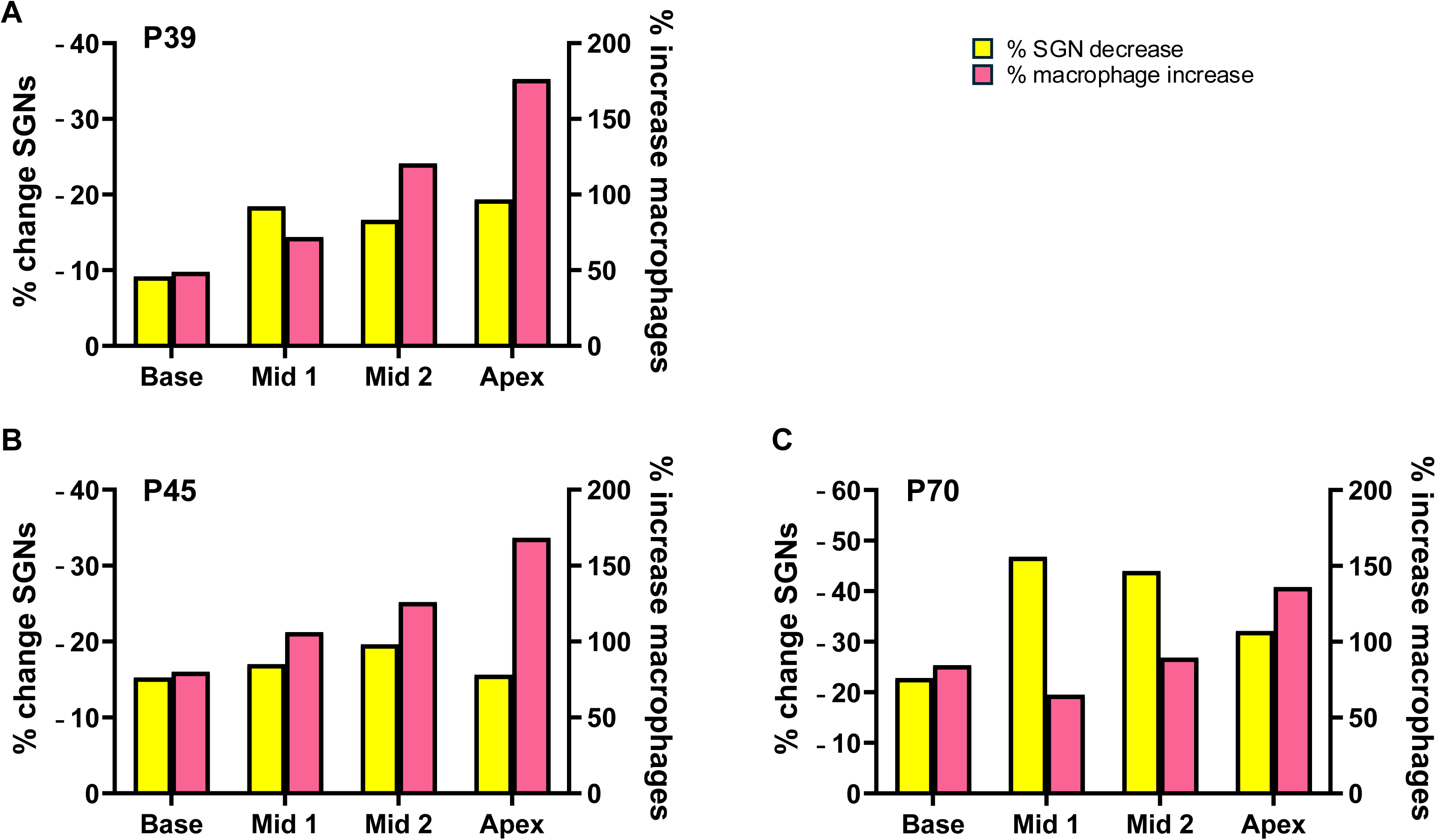
Regional differences in the fractional change (indicated as percent change) of SGN density (decrease) and macrophage density (increase) between hearing and KAN rats at A) P39, B) P45, and C) P70.

